# Sequence-based prediction of protein-protein interactions: a structure-aware interpretable deep learning model

**DOI:** 10.1101/2021.01.22.427866

**Authors:** Samuel Sledzieski, Rohit Singh, Lenore Cowen, Bonnie Berger

## Abstract

Protein-protein interaction (PPI) networks have proven to be a valuable tool in systems biology to facilitate the discovery and understanding of protein function. Unfortunately, experimental PPI data remains sparse in most model organisms and even more so in other species. Existing methods for computational prediction of PPIs seek to address this limitation, and while they perform well when sufficient within-species training data is available, they generalize poorly to new species or often require specific types and sizes of training data that may not be available in the species of interest. We therefore present D-SCRIPT, a deep learning method for predicting a physical interaction between two proteins given just their sequences. Compared to existing methods, D-SCRIPT generalizes better to new species and is robust to limitations in training data size. Our approach encodes the intuition that for two proteins to physically interact, a subset of amino acids from each protein should be in contact with the other. The intermediate stages of D-SCRIPT directly implement this intuition; the penultimate stage in D-SCRIPT is a rough estimate of the inter-protein contact map of the protein dimer. This structurally-motivated design enables interpretability of our model and, since structure is more conserved evolutionarily than sequence, improves generalizability across species. We show that a D-SCRIPT model trained on 38,345 human PPIs enables significantly improved functional characterization of fly proteins compared to the state-of-the-art approach. Evaluating the same D-SCRIPT model on protein complexes with known 3-D structure, we find that the inter-protein contact map output by D-SCRIPT has significant overlap with the ground truth. Our work suggests that recent advances in deep learning language modeling of protein structure can be leveraged for protein interaction prediction from sequence. D-SCRIPT is available at http://dscript.csail.mit.edu.

## 1 Introduction

The systematic mapping of physical protein-protein interactions (PPIs) in the cell has proven extremely valuable in deepening our understanding of protein function and biology. In species such as yeast and humans where a large network of experimentally determined PPIs exists, this PPI network information has proven valuable for downstream inference tasks in understanding functional genomics and biological pathway analysis [1–5]. However, despite the introduction of high-throughput methods [6–10] to assay PPIs, the experimentally determined human PPIs to date are believed to represent only a small fraction of the true set of protein pairs that physically bind in the human cell [11]. In other species, the data is even sparser. As Figure 1 shows, in the case of most model organisms, the number of experimentally determined PPIs is far smaller than for human, and in the case of non-model organisms, it can be nearly non-existent. This motivates our study of computational methods to predict PPIs.

**Figure 1:**
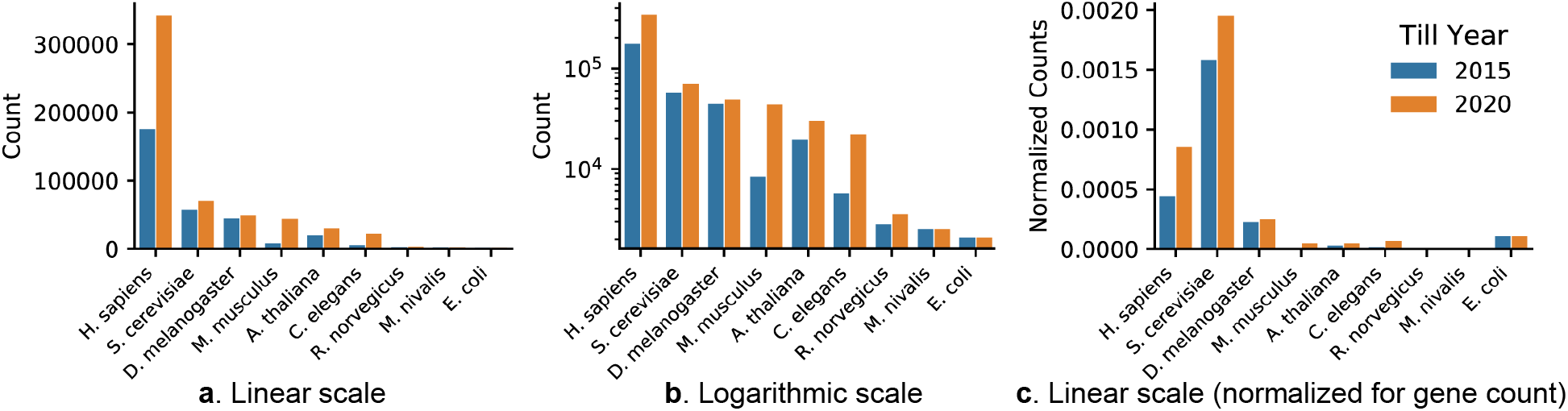
The corpus of experimental PPI data is limited. We show the counts of PPIs in various organisms detected via direct binding experiments (sourced from BioGRID, see Appendix A.1 for details). For readability, we offer both (**a**) linear and (**b**) log-scale plots, as well as (**c**) PPI counts normalized by the square of the species genecount, to adjust for the number of possible gene pairs. Note the relative lack of PPI data in species other than human and yeast. Furthermore, the growth of this corpus over the past five years has been modest (except in the case of human and mouse PPIs).

Here, we introduce a new deep learning method, D-SCRIPT (Deep Sequence Contact Residue Interaction Prediction Transfer), for determining if two proteins interact physically in the cell based on their amino acid sequences. Our key conceptual advance is that a well-matched combination of input featurization and model architecture allow for the model to be trained solely from sequence data, supervised only with a binary interaction label, and yet produce an intermediate representation that substantially captures the structural mechanism of interaction between the protein pair. Our feature construction takes advantage of recent advances in deep learning-based language modeling of single protein structures. Using Bepler and Berger’s [12] pre-trained language model, we construct informative protein embeddings that are endowed with structural information about each of the proteins. The internal representation of our model uses these features to explicitly encode the intuition that a physical interaction between two proteins requires that a subset of the residues in each protein be in contact with the other protein. We do so by having an interprotein contact-map matrix as the penultimate stage in our model, which is then summarized into a single score (Figure 2). These innovations are combined with a novel pooling operation and a regularization loss function that successfully guide the model towards structurally-plausible contact maps during training.

**Figure 2:**
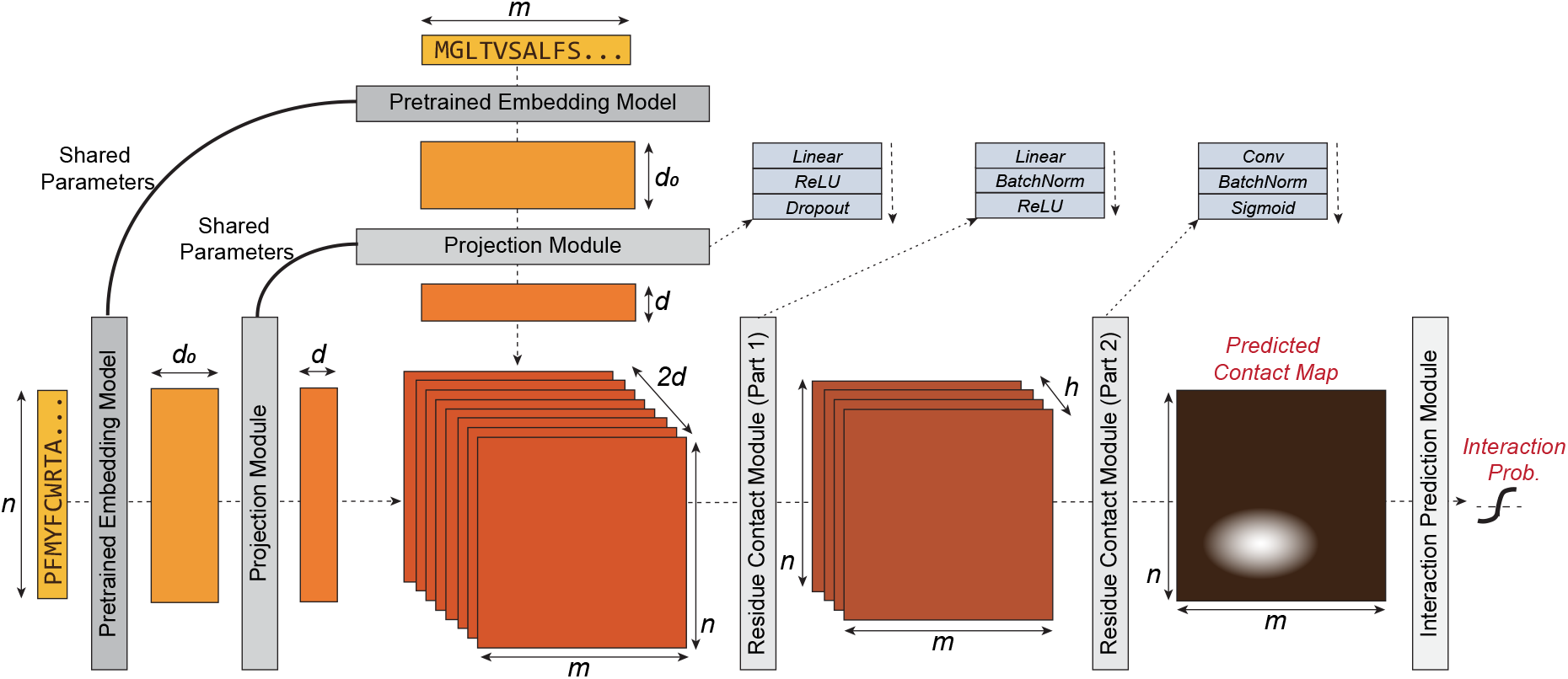
D-SCRIPT architecture overview. Left to right: The Pre-trained Embedding Model generates features for each individual protein. The Projection Model reduces them to *d* dimensions. Each low-dimensional single-protein embedding implicitly includes, among other features, an encoding that broadly captures the protein’s residue-contact map. The novel portion of the architecture combines these low-dimensional embeddings to compute a sparse *interprotein* contact map through a two-step process which first computes a representation for each pair of residues, then incorporates local information using a convolutional filter. The last module (Interaction Prediction Module) uses a customized max-pooling operation (Section 2.1.4) to predict the probability of interaction between the input proteins.

Our design enables D-SCRIPT to offer a combination of advantages that have hitherto been unavailable simultaneously: broad applicability, interpretability, and high cross-species accuracy. It can be applied to arbitrary pairs of protein sequences, including between-species interactions. The intermediate contact-map representation offers a physical interpretation of the interaction. Since structure is more conserved than sequence over evolutionary time [13], this physical model of interaction should translate well across species, making D-SCRIPT more generalizable. We note that the use of Bepler and Berger’s pre-trained model allows us to indirectly benefit from the rich data on 3-D structures of individual proteins. In contrast, a PPI prediction method that was directly supervised with 3-D structures of protein complexes, in order to learn the physical mechanism of interaction, would need to contend with the relatively small size of that corpus [14–16].

We demonstrate D-SCRIPT’s success compared to competing methods in correctly predicting PPIs in organisms with sparse PPI networks. We evaluate D-SCRIPT in the cross-species prediction setting, where a method trained on human PPIs is used to predict PPIs in several less-studied model organisms. We show that in this context, D-SCRIPT substantially improves upon existing methods, including the state-of-the-art deep learning method PIPR [17]. To investigate intra-species performance, we compare D-SCRIPT’s prediction of PPI interactions in the human PPI network, when trained through cross-validation on human PPI interaction data. We find, as expected, that state-of-the-art PIPR substantially outperforms D-SCRIPT when predicting interactions between proteins that have many PPI examples in the training set, but the situation is reversed for proteins with a paucity of PPI interactions in the training set. A simple hybrid method that jointly incorporates the confidence of each method performs best of all.

To examine if the higher cross-species accuracy of D-SCRIPT translates to improved inference of protein function, we run an analysis pipeline to predict functional modules in *Drosophila* using networks predicted by D-SCRIPT and PIPR, and show that D-SCRIPT achieves more functionally coherent clusters. On evaluating the physical plausibility of the intermediate contact map representation, we remarkably find that the map partially discovers the structural mechanism of an interaction despite the model having been trained only on sequence data. Specifically, we evaluate our predictions on Vreven et al.’s [18] benchmark database of 3-D structures of docked protein-pairs and observe that our model’s predicted contact map is significantly similar to the ground-truth inter-protein contact map in cases where our model predicts an interaction. These contact map predictions could be used to augment methods which predict protein interaction from known contacts, such as Hosur et al. [19]. Furthermore, our modular design enables the investigation of model output at various intermediate stages; we demonstrate that each additional stage captures incremental biological intuition. We also present preliminary predictions for viral-host PPIs between SARS-CoV-2 & Human.

### Related Work

D-SCRIPT, like other recent successful deep learning methods PIPR and DPPI [17, 20], belongs to the class of methods that perform PPI prediction from protein amino acid sequence alone, in contrast to a different class of highly successful PPI prediction methods based on network information. Network-based PPI prediction methods use “guilt by association” or more sophisticated network propagation or diffusion methods [4] to computationally predict pairs of proteins that physically interact by bootstrapping from the connectivity patterns of the known interactions [21–24]. Alternatively, other types of protein-protein association that can be more widely available, such as co-expression and co-localization, have also been shown to correlate with physical protein-protein interaction. In cases where there is a reasonably-sized training set and/or a rich set of input data modalities, these methods perform very well and can be powerful tools [3, 25–27]. Unfortunately, there has been limited success in adopting these methods beyond those few model organisms where there is sufficient known interaction (e.g. yeast and Human networks) or association data (e.g yeast networks). Taking advantage of the abundant genetic-interaction coverage in yeast, Khurana et al. [28] described one of the few successful approaches to transfer them to human (where the data is sparse) to glean insights into neurodegenerative disease.

The advantage of a sequence-based approach like D-SCRIPT is that the input sequence data is almost always available, due to the enormous advances in low-cost genome sequencing. Among sequence-based methods, D-SCRIPT’s strength is in its greater cross-species generalizability and more accurate predictions in cases where the existing training data is sparse. Moreover, our contact-map based approach parallels the recent work on deep learning prediction of single-structure contact maps from protein sequence [29–31]. While these methods are trained on 3-D structure data, our method is designed to be trained using only sequence data. Nonetheless, insights from our approach and these methods could potentially be combined in future work.

## 2 Methods

### 2.1 Model Architecture

Our deep learning model for predicting PPIs directly from protein sequences, similar to previous deep learning methods DPPI [20] and PIPR [17], can be composed into two stages:

- Stage 1: Generate a rich feature representation for each protein separately
- Stage 2: Predict an interaction based on these features

where we note that the model is trained end-to-end across both stages. In both DPPI and PIPR, most of the complexity in the model is built into Stage 1. Once a feature representation has been computed, the interaction prediction for DPPI or PIPR is based on a direct element-wise product of the two protein feature vectors. A key innovation of D-SCRIPT is in our design of a more structurally-aware Stage 2. Stage 1 is accomplished by using the pre-trained protein sequence model from Bepler & Berger followed by a projection module (Section 2.1.2), where the model learns low-dimensional protein embeddings which can also be used as compact representations for downstream interaction and structural prediction tasks. For the second stage, we present a novel architecture that encodes a *physical* model of protein interaction: we predict two proteins to interact only if there exists a short sequence of residues in the first protein that is highly compatible with a sequence of residues in the second protein. In the contact module (Section 2.1.3), the low-dimensional embeddings are used to compute a sparse contact map which predicts the locations of contacts between protein residues. Finally, the interaction module (Section 2.1.4) uses a customized max-pooling operation on the contact map to predict the probability of interaction between the input proteins.

#### 2.1.1 Input &Structure-aware Embedding

The input to D-SCRIPT is a pair of protein sequences *S*_1_, *S*_2_ with lengths (*n, m*) and it outputs an interaction probability 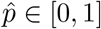 and a predicted-contact matrix *Ĉ ∈* [0, 1]^*n×m*^.

We first generate embeddings 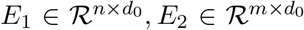 by embedding *S*_1_ and *S*_2_ with a pre-trained model from Bepler & Berger. Their model is a Bi-LSTM (bidirectional long short-term memory) neural network trained on three independent pieces of information: 1) the protein’s SCOP classification, indicating its general structure, 2) self-contact map of a protein’s 3-D structure, and 3) sequence alignment of similar proteins. These embeddings capture both local and global structural features of the protein sequences: the *d*_0_-dimensional encoding of each amino acid contains information not just about the amino acid and its immediate context, but also the global structure of the protein. This is a key distinction from other approaches (e.g. Chen et al.’s in PIPR), where each amino acid’s embedding represents just its biochemical properties or a short-range context (e.g., 5-7 residues) around it. We note that alternative embeddings (e.g. Rives et al. [32] or Luo et al. [33]) can potentially be substituted here.

#### 2.1.2 Projection Module

In the projection module, we reduce *E*_1_ and *E*_2_ to *d*-dimensional representations using a fully-connected linear layer (multi-layer perceptron) with *d*_0_ input and *d* output nodes. Specifically, given an input embedding 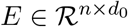, we compute the embedding projection 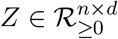 as

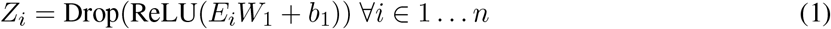

with 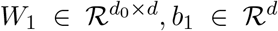 as learned weights and biases. The rectified linear unit (ReLU) is a non-linear operation which applies the transformation ReLU(*x*) = max(0, *x*). The dropout layer (Drop) randomly sets 50% of the weights to zero, helping prevent over-fitting in *W*_1_.

#### 2.1.3 Residue Contact Module

The residue contact model takes the *d*-dimensional embeddings *Z*_1_, *Z*_2_ and models the interaction between the residues of each protein. First, for each pair of residue embeddings 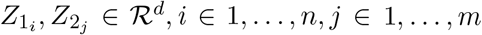, we compute a broadcast matrix with hidden dimension *h*, 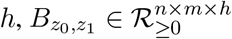

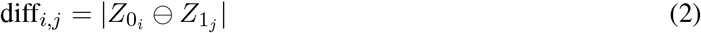

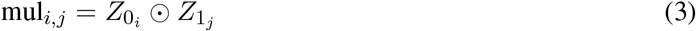

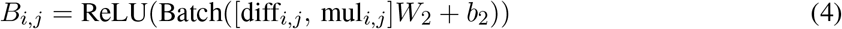

where ⊖ indicates the element-wise difference and ⊙ indicates the Hadamard product. This featurization is symmetric and has been previously used in NLP and protein sequence modeling tasks [12, 34].

*W*_2_ *∈ℛ*^2*d×h*^, *b*_2_ *∈ℛ*^*h*^ are the learned weights and biases. The batch normalization operation normalizes the mean and variance of the input features, thus stabilizing the learning process. Each element *B*_*i,j*_ captures the direct interaction between residues 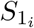 and 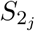. The broadcast matrix *B* is used to compute the contact prediction matrix *Ĉ ∈* [0, 1]^*n×m*^, where

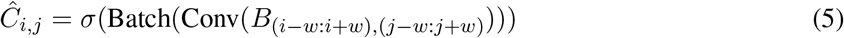

The two-dimensional convolution (Conv) operation with width 2*w* + 1 and *h* channels uses the *h*-dimensional representation of all residues within *w* of *B*_*i,j*_ to compute *Ĉ*_*i,j*_, and thus detects local patterns in two-dimensional residue contact space. The broadcast matrix is zero-padded to allow for the convolution operation to be performed at all indices. We again apply a batch normalization to stabilize learning. We apply the sigmoid operation *σ*, which restricts the output values of *Ĉ* to be in the range [0,1], and thus they can be interpreted as the predicted probability that two residues are in contact.

#### 2.1.4 Interaction Prediction Module

The interaction prediction module computes a single probability of interaction 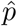 from the *n×m* contact map *Ĉ*. To do so, we perform two pooling operations. The first is a standard max-pool: an *l*-dimensional max-pool divides *Ĉ* into 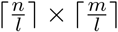 non-overlapping regions and takes the maximum value of each region, with zero-padding applied where necessary. This max-pooled matrix *P* represents the probability of interaction in local regions of the proteins and maintains only the highest-probability residue contacts in each region for global prediction. The second pooling operation is a global pooling operation, calculated as

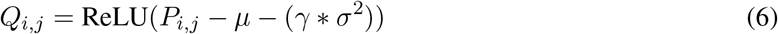

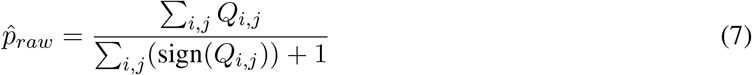

where *µ, σ*^2^ are the mean and variance of the *P*_*ij*_ values and *γ* is a learned parameter. The matrix *Q* sparsifies *P*, maintaining only those contacts which are *γσ*^2^ greater than the mean, and setting all others to zero. We then predict that the proteins will interact with the average probability of interaction among these high-probability contacts. Together with the regularization that the contact matrix be sparse (see Section 2.2), this global pooling operation captures the intuition that a pair of interacting proteins will be characterized by a relatively small number of high-probability interacting residues or regions.

The final step of interaction prediction is designed to enhance the bimodality of the output distribution, so that the choice of a cutoff becomes less important in distinguishing positive and negative predictions. We apply the logistic activation function to compute the output probability 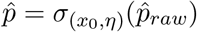 where

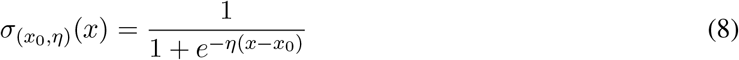

This activation function, with *x*_0_ = 0.5, takes our raw probability predictions and makes them more “extreme”, depressing values below *x*_0_ towards 0 and inflating values above *x*_0_ towards 1, with *η* controlling the rate at which this occurs. We return 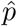 and *Ĉ* as the model prediction, from which we calculate the loss and optimize the gradient as described in Section 2.2.

### 2.2 Training

#### Training Objective

Given the true labels, the predicted probabilities 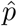, and the contact maps *Ĉ*, we compute the loss as *λL*^*BCE*^ + (1 *− λ*)*L*^*MAG*^; here *λ* is a hyper-parameter that balances between *L*^*BCE*^, the binary cross-entropy (BCE) loss, and *L*^*MAG*^, the contact-map magnitude loss (MAG). While the BCE loss is standard in a classification context, we introduce *L*^*MAG*^ as a novel regularization that enables us to learn realistically sparse contact maps. *L*^*MAG*^ for a single training example is calculated as the arithmetic mean value of the contact map *Ĉ*. Jointly minimizing the total magnitude of contact maps with the BCE captures the intuition that interacting proteins are characterized by just a few high probability inter-protein contacts, while most residues will not be in contact.

#### Implementation Details

We implemented D-SCRIPT in PyTorch 1.2.0 and trained with a NVIDIA Tesla V100 with 32GB of memory. Embeddings from the pre-trained Bepler & Berger model were produced by concatenating the final values of the output and all hidden layers, so that *d*_0_ = 6165. We used a projection dimension of *d* = 100, a hidden dimension of *h* = 50, a convolutional filter with width 2*w* + 1 = 7, and a local max-pooling width of *l* = 9. We used *x*_0_ = 0.5, *η* = 20 for the custom logistic activation, and *λ* = 0.35 for calculating the training loss. Weights were initialized using PyTorch defaults. We used a batch size of 25, the Adam optimizer with a learning rate of 0.001, and trained all models for 10 epochs.

## 3 Results

### PPI Data Set

To evaluate the performance of D-SCRIPT in predicting protein-protein interactions, we use data from the STRING [26] database (version 11). STRING contains protein pairs corresponding to a variety of primary sources and interaction modalities (e.g., binding vs co-expression). In order to select only high-confidence physical protein interactions, we limited our positive examples to binding interactions associated with a positive experimental-evidence score. From this set, we removed PPIs involving very short proteins (shorter than 50 amino acids) and, due to GPU memory constraints, also excluded proteins longer than 800 amino acids. Next, we removed PPIs with high sequence redundancy to other PPIs, following the precedent of previous approaches [17, 20]. Specifically, we clustered proteins at the 40% similarity threshold using CD-HIT [35, 36], and a PPI (A-B) was considered sequence redundant (and excluded) if we had already selected another PPI (C-D) such that the protein pairs (A, C) and (B, D) each shared a CD-HIT cluster. Removing sequence redundant PPIs from the data set prevents the model from memorizing interactions based on sequence similarity alone. To generate negative examples of PPI, we followed [20] and randomly paired proteins from the non-redundant set, choosing a 10:1 negative-to-positive ratio to reflect the intuition that true positive PPIs are likely rare. Our human PPI data set contained 47,932 positive and 479,320 negative protein interactions, of which we set apart 80% (38,345) for training and 20% (9,587) for validation. For each of 5 model organisms (Table 1) we selected 5,000 positive interactions and 50,000 negative interactions using this procedure, with the exception of *E*.*coli* (2,000/20,000) where the available set of positive examples in STRING was limited.

**Table 1:**
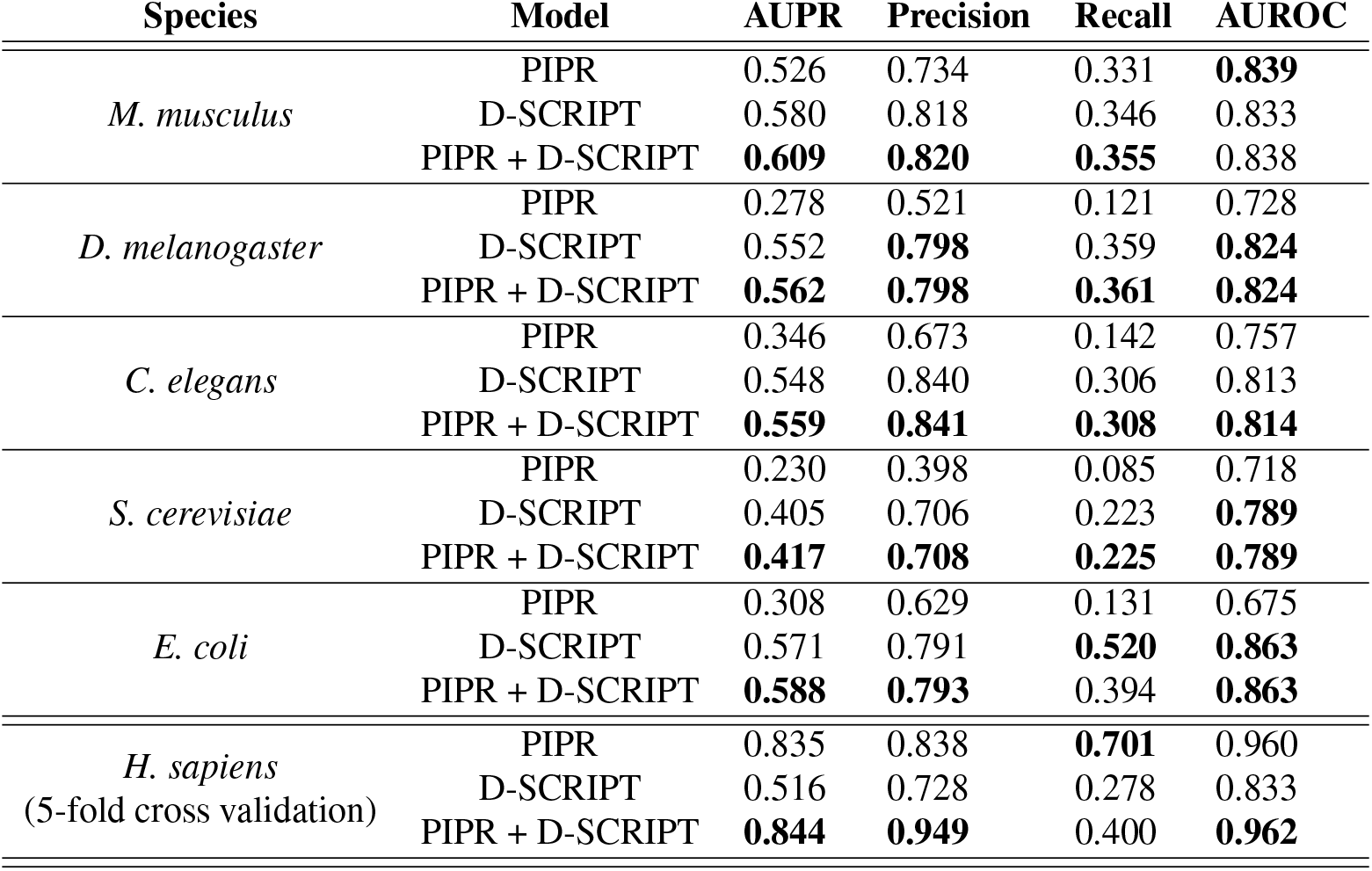
Evaluation of models trained on human PPIs. *H. sapiens* results are average performance over 5-fold cross validation. All other species were evaluated using a model trained on human data. D-SCRIPT significantly outperforms PIPR cross-species, despite PIPR performing better in human cross-validation. In both cases, a hybrid model is able to achieve better performance than either model alone.

| Species | Model | AUPR | Precision | Recall | AUROC |
| --- | --- | --- | --- | --- | --- |
| *M. musculus* | PIPR | 0.526 | 0.734 | 0.331 | **0.839** |
|  | D-SCRIPT | 0.580 | 0.818 | 0.346 | 0.833 |
|  | PIPR + D-SCRIPT | **0.609** | **0.820** | **0.355** | 0.838 |
| *D. melanogaster* | PIPR | 0.278 | 0.521 | 0.121 | 0.728 |
|  | D-SCRIPT | 0.552 | **0.798** | 0.359 | **0.824** |
|  | PIPR + D-SCRIPT | **0.562** | **0.798** | **0.361** | **0.824** |
| *C. elegans* | PIPR | 0.346 | 0.673 | 0.142 | 0.757 |
|  | D-SCRIPT | 0.548 | 0.840 | 0.306 | 0.813 |
|  | PIPR + D-SCRIPT | **0.559** | **0.841** | **0.308** | **0.814** |
| *S. cerevisiae* | PIPR | 0.230 | 0.398 | 0.085 | 0.718 |
|  | D-SCRIPT | 0.405 | 0.706 | 0.223 | **0.789** |
|  | PIPR + D-SCRIPT | **0.417** | **0.708** | **0.225** | **0.789** |
| *E. coli* | PIPR | 0.308 | 0.629 | 0.131 | 0.675 |
|  | D-SCRIPT | 0.571 | 0.791 | **0.520** | **0.863** |
|  | PIPR + D-SCRIPT | **0.588** | **0.793** | 0.394 | **0.863** |
| *H. sapiens*<br>(5-fold cross validation) | PIPR | 0.835 | 0.838 | **0.701** | 0.960 |
|  | D-SCRIPT | 0.516 | 0.728 | 0.278 | 0.833 |
|  | PIPR + D-SCRIPT | **0.844** | **0.949** | 0.400 | **0.962** |

### D-SCRIPT Excels at Cross-Species PPI Prediction

We first sought to see how D-SCRIPT performed on the task of cross-species interaction prediction. We trained a model on human PPIs and evaluated it using PPI data sets from five other model organisms. We compared D-SCRIPT with PIPR, shown by Chen et al. [17] to be the best performing sequence-based PPI prediction method, trained on the same set of human PPIs as the D-SCRIPT model. We compare model complexity of D-SCRIPT and PIPR in Appendix A.2. We also compare to a recently released method by Richoux et al. [37] in Appendix A.3. Additionally, we compared with a hybrid method (PIPR + D-SCRIPT) where PIPR was used to adjust D-SCRIPT’s prediction: when PIPR is highly confident that an interaction *does* occur 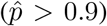, the predicted probability of D-SCRIPT is increased by 50%. In Table 1, we report the precision, recall, area under precision-recall curve (AUPR) and area under ROC curve (AUROC) of each method in each of five species. For highly unbalanced data, as is the case here, we note that AUPR is generally considered a better metric than AUROC. D-SCRIPT significantly outperforms PIPR cross-species and maintains a high AUPR across all species, even those which are extremely evolutionary distant from human. In fact, its AUPR in these species remains comparable to that seen in human cross-validation. The hybrid method outperforms both D-SCRIPT and PIPR alone, but improves D-SCRIPT only modestly in cross-species analysis.

### D-SCRIPT Hybrid Performs Well on Human Cross Validation

While our aim is enhanced cross-species PPI prediction, we sought to investigate how D-SCRIPT would perform at predicting within-species interactions in Human. We performed 5-fold cross validation and report here the average across all folds. Additionally, we evaluated a hybrid method (PIPR + D-SCRIPT) where we used D-SCRIPT to adjust PIPR’s prediction: when D-SCRIPT is highly confident that an interaction does *not* happen 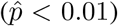, we reduce the predicted probability from PIPR by half. Table 1 shows that while PIPR significantly outperforms D-SCRIPT on human PPIs in cross-validation, a combination of the methods outperforms either one alone. Notably, the PIPR + D-SCRIPT hybrid achieves substantially higher precision, though at the expense of recall. This may be a desirable trade-off in certain contexts, e.g., when generating PPI candidates for experimental validation. We further investigated the performance of D-SCRIPT and PIPR on subsets of the human data, seeking to better understand their relative strengths. As we elaborate in Appendix A.4, D-SCRIPT performs better on interactions involving proteins that occur infrequently in the PPI network while PIPR performs better on proteins that occur frequently. This suggests that PIPR may perform better when a large amount of training data is available but D-SCRIPT may generalize better to new proteins (and species).

### Functional Module Prediction

The importance of PPI networks arises, in part, from the graph-theoretic analyses on them which enable the functional characterization of un-annotated proteins. We therefore sought to test if D-SCRIPT’s success at cross-species generalization would translate to better functional inference in new species. In particular, we hypothesized that, compared to PIPR, the D-SCRIPT model trained on human data should facilitate more accurate inference of protein functional modules in *Drosophila*. Towards this end, we generated a set of 10,475,595 candidate pairs from the set of *Drosophila* proteins in STRING. Using D-SCRIPT and PIPR’s human-trained models, we predicted interactions over this candidate set. Module detection on the resulting PPI networks was performed by spectral clustering, with pairwise distances between proteins assessed using Cao et al.’s [38, 39] diffusion state distance (DSD) metric. This module detection approach performed well in a recent DREAM challenge on functional module detection [5].

We tested the relative functional coherence of 374 (PIPR) and 384 (D-SCRIPT) clusters using available GO (Gene Ontology) annotations from FlyBase [40], filtering out electronically-inferred and homology-based annotations. All GO terms were mapped to a limited set of GO Slim terms using the *D. Melanogaster* species-specific list [41]. For each cluster, we computed the within-cluster functional similarity, calculated as the mean Jaccard similarity of the sets of GO Slim annotations for all pairs of proteins within the cluster. We also used a different, graph-theoretic measure of protein similarity based on GO terms [42]; it produced similar results (Appendix A.5). Figure 3 shows that the average within-cluster similarity when interactions are predicted using D-SCRIPT is significantly higher than when using PIPR (*p* = 0.000723, one-tailed *t*-test), and that D-SCRIPT results in 24% more highly-enriched (top 10% of scores) clusters as PIPR.

**Figure 3:**
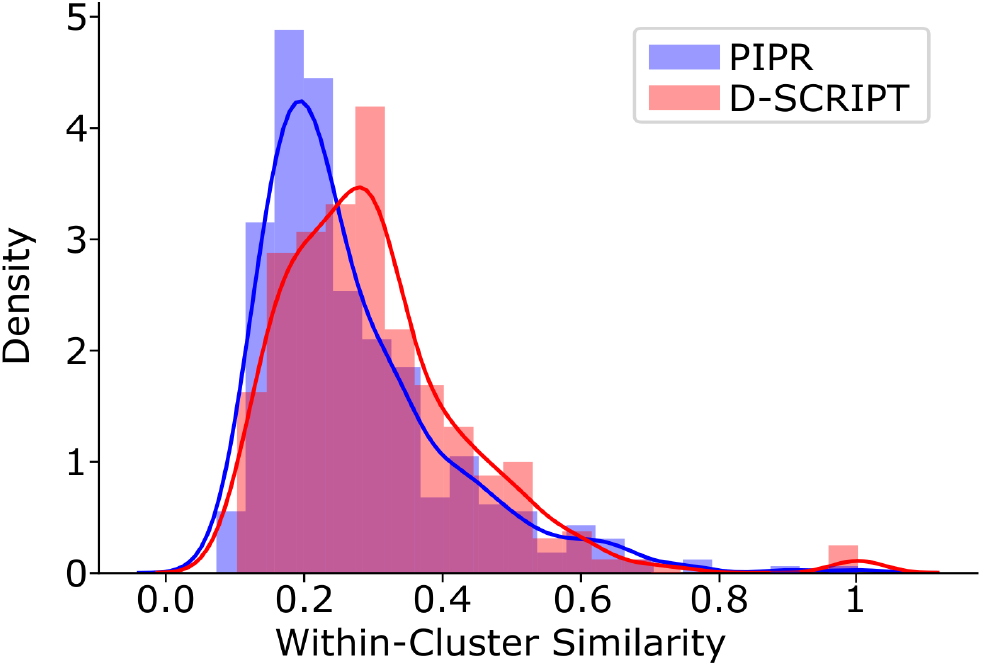
Distribution of within-cluster similarity scores in *Drosophilia melanogaster*. D-SCRIPT recovers more functionally coherent clusters than PIPR (*p* = 0.000723, one-tailed *t*-test). Clusters were generated by predicting protein interactions using either D-SCRIPT or PIPR, computing the diffusion state distance (DSD) between proteins, and clustering the DSD matrix using spectral clustering. Within-cluster similarity was calculated as the average Jaccard similarity between GO Slim annotations of all pairs of proteins in the cluster.

### Preliminary prediction of viral-host PPIs between SARS-Cov-2 & Human

We applied D-SCRIPT and PIPR to computationally screen all SARS-Cov-2 proteins against 19,777 human proteins, predicting approximately 3,000 viral-host PPIs from each method (Appendix A.6). We then characterized each viral protein’s function by the GO annotations of its human interactors. Compared to the corresponding annotations derived from 332 experimentally-determined PPIs (Gordon et al. [43]), we found D-SCRIPT based annotations to overlap more with the experimental results than those from PIPR (*p* = 0.059, paired one-tailed *t*-test). We have made these predicted PPIs available for the community’s use (see A.6), but note that these are preliminary results and need further analysis.

### Comparison of D-SCRIPT Embedding with Other Embeddings

To compare our sequence embedding method with other potential ones, we evaluated various protein sequence embeddings under the following framework: the Euclidean distance in an embedding space was used to define a distance measure between proteins. Given a true positive PPI (*A, B*), we applied this distance measure to identify the *k*-nearest neighbors of *A* and *B* each, and computed how many of the *k*^2^ possible combinations of these neighbors corresponded to a positive PPI. For D-SCRIPT, we used the 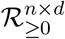 output of the projection module (Section 2.1.2) and averaged the features across the length of the protein to obtain a *d*-dimensional embedding. We compared with a one-hot embedding categorizing each amino acid into one of seven classes based on biochemical properties (“AAClass”, from [44]), a 5-residue-context Skip-Gram embedding (“Vec5”, from [45]), a concatenation of Vec5 and AAClass (used in the input for PIPR), and a randomly generated 50-dimensional embedding with values drawn uniformly from the range [0, 1]. Additionally, we used BLAST [46] to select *k* neighbors. Figure 4 shows that D-SCRIPT significantly outperforms all other embeddings.

**Figure 4:**
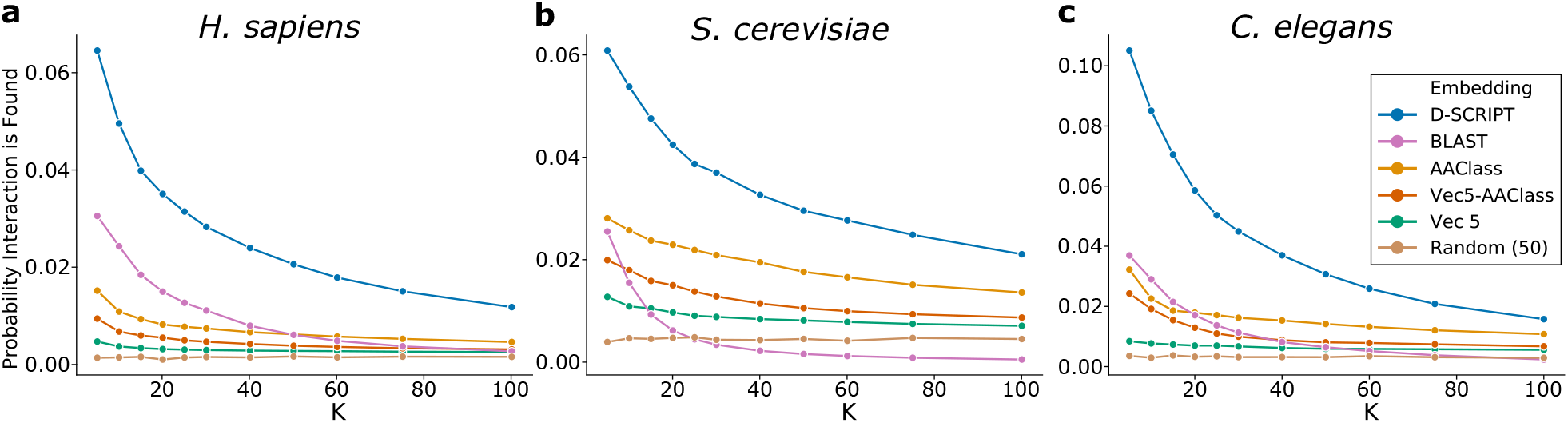
Neighborhood interaction search using protein embeddings. Performance comparison of D-SCRIPT projection module (Section 2.1.2) trained on human data with several other embedding methods on recovering true interacting protein pairs in the neighborhood of known PPIs in human (**a**), yeast (**b**), and roundworm (**c**). D-SCRIPT embeddings recover significantly more interacting proteins than any other embedding, regardless of species or number of neighbors checked. AAClass also performs well, likely because it characterizes biochemistry which is preserved at longer evolutionary distances. BLAST performs well at low values of *k* but has difficulty recovering interactions for larger values — likely due to network rewiring over longer evolutionary distances.

### Prediction of Intra-Protein Self Contacts

One of our aims when designing D-SCRIPT was to capture the structural aspects of interaction — the per-protein embedding produced by the trained projection module (Section 2.1.2) should encode structural information. To examine this aspect, we randomly selected 300 proteins from the Protein Data Bank (PDB) and generated (*n× d*)-dimensional embeddings of these proteins using the D-SCRIPT model trained on human PPIs. We assessed intra-protein contacts at 8 Å in the PDB structure and converted each protein’s contact map to a binary classification data set: a protein sequence of length *n* corresponded to 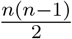 observations, with the observation *ij* corresponding to a putative contact between residues *i* and *j*. We randomly split the 300 PDB structures into a training set of 100 and a test set of 200, evaluating how well a logistic regression that uses the D-SCRIPT embeddings as the input predicts contacts between residues. The regression was *L*_2_-regularized and class balanced, with its input for observation *ij* being the concatenation of the *d*-dimensional embeddings *Z*_*i*_ and *Z*_*j*_ output by the projection module as well as their combinations diff_*ij*_, mul_*ij*_ as defined in Equations 2 & 3. Figure 5 shows that a linear combination of features in the projection module output is able to recapitulate a significant subset of the ground-truth contacts, achieving a median per-structure AUPR of 0.19 over the test data set. These results strongly suggest that the end-to-end training of D-SCRIPT — using only sequence data — results in an intermediate representation that captures structural information at the level of each protein.

**Figure 5:**
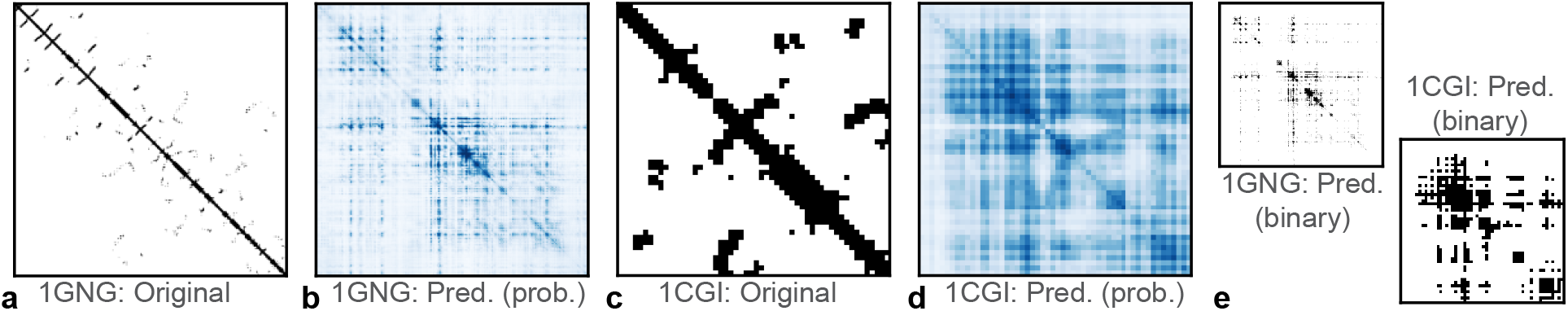
Self-contact prediction using D-SCRIPT embeddings. The PDB identifier 1GNG corresponds to a protein with 356 residues where the D-SCRIPT embedding’s self-contact prediction accuracy is near the median of cases we studied (AUPR=0.19), while 1CGI corresponds to a short protein (54 residues) in which the embedding achieves a higher accuracy (AUPR=0.38). The original contacts (**a, c**) were assessed at 8 Å. Contacts were predicted (**b, d**) using a logistic regression model trained on contacts seen in a held-out set of proteins, and the binarization thresholds for panel (**e**) were chosen so as to result in the same number of contacts as in the original maps.

### Prediction of Inter-Protein Docking Contacts

We investigated whether the interpretability of our model could aid in predicting inter-protein docking contacts. The output of the Residue Contact Module (Section 2.1.3) of D-SCRIPT is an inter-protein contact map *Ĉ* where *Ĉ*_*ij*_ *∈*[0, 1] can be interpreted as the probability of residue *i* from protein *S*_1_ being in contact with residue *j* of protein *S*_2_. We verified the contact maps produced after training were consistent with our design goal: the maps corresponding to negative predictions should have uniformly near-zero *Ĉ*_*ij*_ scores, while those for positive predictions should be sparse but with isolated regions of high *Ĉ*_*ij*_ scores. We found that this was indeed the case generally and show some examples in Figure 6: the maximum *Ĉ* value is high for positive examples and low for negative examples.

**Figure 6:**
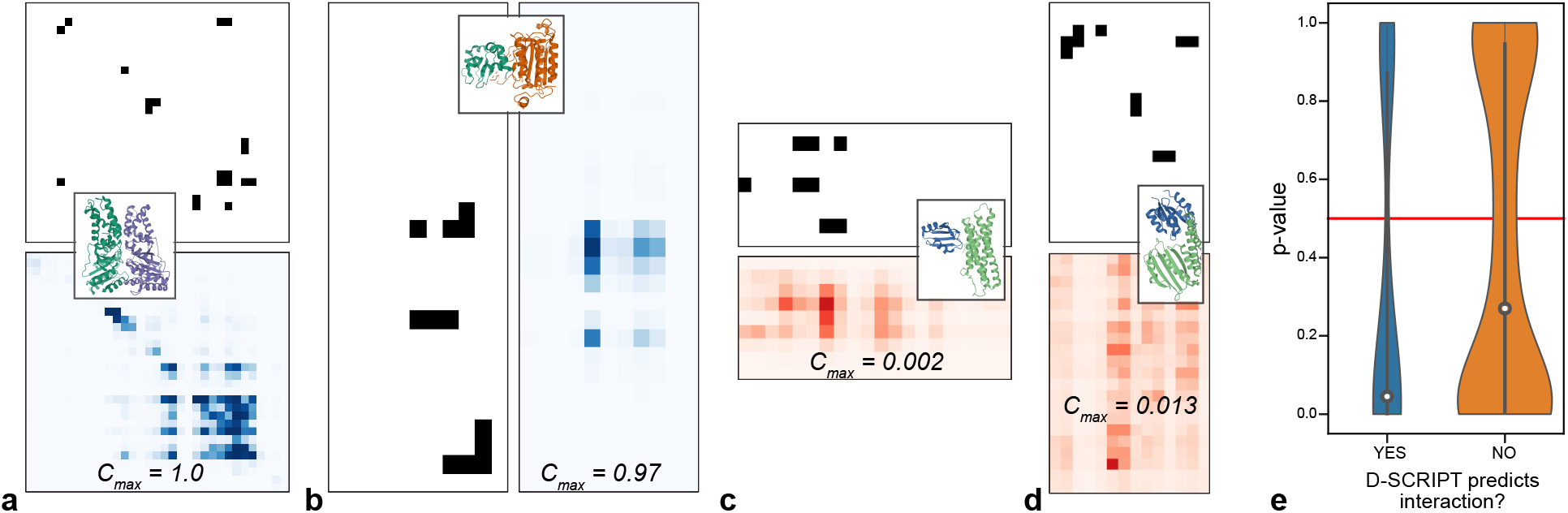
When D-SCRIPT correctly predicts an interaction, its contact maps *Ĉ*are significantly similar to the ground truth. We show inter-protein contact maps of protein structures known to dock together. Panels **(a**,**b)** correspond to pairs where D-SCRIPT correctly predicted an interaction, while panels **(c**,**d)** are cases where it incorrectly predicted no interaction. The black-and-white matrices correspond to the PDB ground truth while the colored matrices correspond to D-SCRIPT’s *Ĉ*; for the latter, the color scales of (a,b) differ from (c,d). While *Ĉ* contains large values for positive pairs, its maximum *C*_*max*_ is very low for negative pairs. Panel **(e)** shows a systematic evaluation of the 2-D Earth mover’s distance-based similarity between *Ĉ* and the ground truth. Not only are the correctly-predicted *Ĉ*s significantly similar to the ground truth, even when D-SCRIPT incorrectly predicts two proteins don’t interact, its contact maps are still similar to ground truth.

We next sought to test if *Ĉ* is physically representative of the actual docking mechanism of the interaction. We emphasize that this is a high bar given we do not provide any 3-D information to the model nor any guidance on docking and, in principle, the model could perform well on the classification task without *Ĉ* being physically accurate. We performed this test using Hwang et al.’s benchmark data set of docked protein structures [18]. For every pair of chains in each PDB complex in the benchmark set, we generated a candidate PPI. We applied our human-data-trained model on 295 candidate PPIs and evaluated the predicted contact maps against the ground-truth contacts (assessed at 8 Å). In cases where our model predicted an interaction, we found *Ĉ* to indeed recapitulate the ground-truth contacts substantially (Figure 6 a,b). Even in some of the cases where D-SCRIPT did not predict an interaction, the distribution of *Ĉ*_*ij*_ scores was nevertheless consistent with the ground-truth (Figure 6 c). To systematically evaluate *Ĉ*’s accuracy, we interpreted ground truth and predicted contact maps as probability distributions over the *n ×m* matrix and measured the 2-D Earth mover’s distance between these distributions, computed by solving an optimal transport problem under the Euclidean metric [47]. We chose this metric to measure similarity between regions of the two contact maps rather than measuring per-residue matches (with a metric such as binary cross-entropy) because the convolutional and max-pooling layers in our model aggregate over neighboring residues, thus diffusing the signal. For each candidate PPI, we established random baselines by shuffling *Ĉ* and recomputing the Earth mover’s distance. We estimated the p-value of the actual *Ĉ* against 500 random trials, finding that in cases where D-SCRIPT predicted an interaction, the *Ĉ* contact-maps were significantly similar to the ground-truth (median FDR-corrected *p* = 0.08). Even in cases where D-SCRIPT did not predict an interaction, *Ĉ*’s similarity to the ground truth was higher than that of the random baselines.

### Performance

D-SCRIPT took approximately 3 days to train for 10 epochs on 843,602 training pairs, and fits within a single 32GB GPU. Running time and GPU memory usage scales roughly quadratically, O(*mn*), with the protein lengths *m, n*, since D-SCRIPT models the full *n ×m* contact map as an intermediate step. Prediction of new candidate pairs with a trained model is very fast, requiring on average 0.13 seconds/pair and less than 5GB of GPU memory. Since D-SCRIPT generalizes well cross-species, it only needs to be trained once on a large corpus of data, and can be used to make predictions in a variety of settings.

## 4 Discussion

We have introduced D-SCRIPT, an interpretable method for PPI interaction prediction from sequence. It joins the small but growing set of advances in interpretable deep learning methods in computational biology [48, 49]. We showed that its predictions generalize better than other methods to PPIs for which there are fewer training examples of interactions with the constituent proteins, and importantly, to a cross-species setting where the model is only trained on protein sequences from a different species. This is particularly exciting for organisms such as *C. elegans*, for which the known portion of the PPI network is still quite sparse. In organisms like *D. melanogaster* where some PPI data does exist, future work could productively combine that data with D-SCRIPT models trained on human data. We suspect that D-SCRIPT’s relative success across species, but under-performance on a within-species evaluation, is due to the simplicity of the model and the extent to which it is regularized. These design choices enhance D-SCRIPT’s generalizability, directing it to learn general structural aspects of the interaction, rather than using network structure or the frequency of any individual protein as an interaction partner. However, for certain tasks a balance between the cross-species generalizability of D-SCRIPT and the within-species specificity of other state-of-the-art methods may be desirable. A future research direction might be transfer learning to tune a pre-trained D-SCRIPT model to a target species, while another approach could be to integrate it with graph-theoretic PPI predictions. Insights from recent advances in the prediction of contact maps and structures of individual proteins could also be incorporated into our model architecture. D-SCRIPT illustrates that learning the language of individual proteins, a remarkably successful deep learning effort, also helps decode the language of protein interactions.

## Acknowledgements

We thank Tristan Bepler for helpful discussions and technical assistance. SS, RS and BB were supported by the NIH grant R01 GM081871. LC was supported by NSF HDR grant 1939263.

## A Appendix

### A.1 The Corpus of Experimental PPI Data is Limited

We sourced the PPI data in Figure 1 from BioGRID. While we have sourced PPI data from the STRING database everywhere else in this paper, here we chose to use BioGRID because the publication date of a PPI is easily accessible in BioGRID, allowing us to estimate the number of PPIs assayed in the last five years. While the BioGRID selection may not precisely match the STRING selection due to curation differences between the two databases, our primary aim here is conveying the relative data availability across species; this estimate should not be significantly impacted by differences in curation.

### A.2 Comparison of Model Complexities of D-SCRIPT and PIPR

The version of PIPR that we compare to has 72,500 trainable parameters. The full D-SCRIPT model has 629,207 trainable parameters, but the vast majority of those (616,600) are in the projection module, which is a simple linear combination of all concatenated hidden states of the Bepler & Berger language model (which are themselves already redundant). The remaining stages of the model together have 12,707 trainable parameters. Practically, we prevent over-fitting through a high rate of dropout (50%) in the Projection module (Section 2.1.2), combined with a very simple but structurally informed architecture in the Contact (Section 2.1.3) and Interaction (Section 2.1.4) prediction modules. Additionally, the contact map magnitude loss (Section 2.2) acts as a regularization by requiring that the predicted contact maps be sparse. Empirically, D-SCRIPT generalizes much better than PIPR, and does not seem to overfit to the training data.

### A.3 Comparison with Another Sequence-based PPI Prediction Method

We also compared our method to a model described by Richoux et al. [37]. In it, they trained a multi-layer fully connected network to predict protein interaction given two input sequences. The results comparing this model to D-SCRIPT, presented in Table A.1, are organized like those presented in Table 1 in the main text. Similarly to Chen et al.’s PIPR model, the Richoux et al. model performs better than D-SCRIPT when trained and tested on human PPIs while D-SCRIPT generalizes better across species, with substantially higher AUPRs than the Richoux model when the human-data-trained model is evaluated on other species. “Richoux + D-SCRIPT” refers to the same hybrid combination described in Section 3.

### A.4 Deeper Investigation of D-SCRIPT and PIPR’s Relative Strengths

We investigated the relative performance of D-SCRIPT and PIPR on subsets of human proteins, seeking to better understand their relative strengths.

#### Using protein frequency counts as a predictor

PPI networks are dominated by a dense and highly connected core of nodes, where the core includes many hub nodes. A method that does relatively well on these hub proteins will have an advantage on within-species evaluations. For our cross-validation setup, we followed the precedent of previous work and split training and test data by edges [17, 20]. Consequently, a protein’s relative frequency will be roughly similar in the training and test data sets. In this context, a PPI prediction heuristic that simply remembers frequency counts does strikingly well (see figure below). Here, we scored the likelihood of a candidate PPI (*A, B*) being true as the minimum of frequency counts, in training data, of *A* and *B*.

**Table A.1:** Evaluation of models trained on human PPIs. We compare D-SCRIPT to the fully connected model described in Richoux et al. by a 5-fold cross validation in human, and the application of a human-data trained model to PPIs from 5 other species. As with the comparison to PIPR (see Table 1), while Richoux outperforms PIPR in the cross validation setting, D-SCRIPT generalizes significantly better to unseen proteins cross-species, and a hybrid model outperforms either model alone.

| Species | Model | Precision | Recall | AUPR | AUROC |
| --- | --- | --- | --- | --- | --- |
| *M. musculus* | Richoux | 0.749 | 0.321 | 0.518 | **0.816** |
|  | D-SCRIPT | 0.818 | 0.346 | 0.580 | 0.833 |
|  | Richoux + D-SCRIPT | **0.821** | **0.354** | **0.608** | 0.837 |
| *D. melanogaster* | Richoux | 0.582 | 0.126 | 0.231 | 0.659 |
|  | D-SCRIPT | 0.798 | 0.359 | 0.552 | 0.824 |
|  | Richoux + D-SCRIPT | **0.799** | **0.360** | **0.559** | **0.824** |
| *C. elegans* | Richoux | 0.672 | 0.079 | 0.252 | 0.671 |
|  | D-SCRIPT | 0.840 | 0.306 | 0.548 | 0.813 |
|  | Richoux + D-SCRIPT | **0.841** | **0.307** | **0.554** | **0.814** |
| *S. cerevisiae* | Richoux | 0.501 | 0.059 | 0.201 | 0.652 |
|  | D-SCRIPT | 0.706 | 0.223 | 0.405 | 0.789 |
|  | Richoux + D-SCRIPT | **0.707** | **0.224** | **0.413** | **0.789** |
| *E. coli* | Richoux | 0.746 | 0.093 | 0.303 | 0.687 |
|  | D-SCRIPT | 0.791 | 0.520 | 0.571 | 0.863 |
|  | Richoux + D-SCRIPT | **0.792** | **0.391** | **0.581** | **0.863** |
| *H. sapiens*<br>(5-fold cross validation) | Richoux | 0.815 | **0.737** | 0.834 | 0.964 |
|  | D-SCRIPT | 0.728 | 0.278 | 0.516 | 0.833 |
|  | Richoux + D-SCRIPT | **0.944** | 0.405 | **0.848** | **0.966** |

**Figure A.1:**
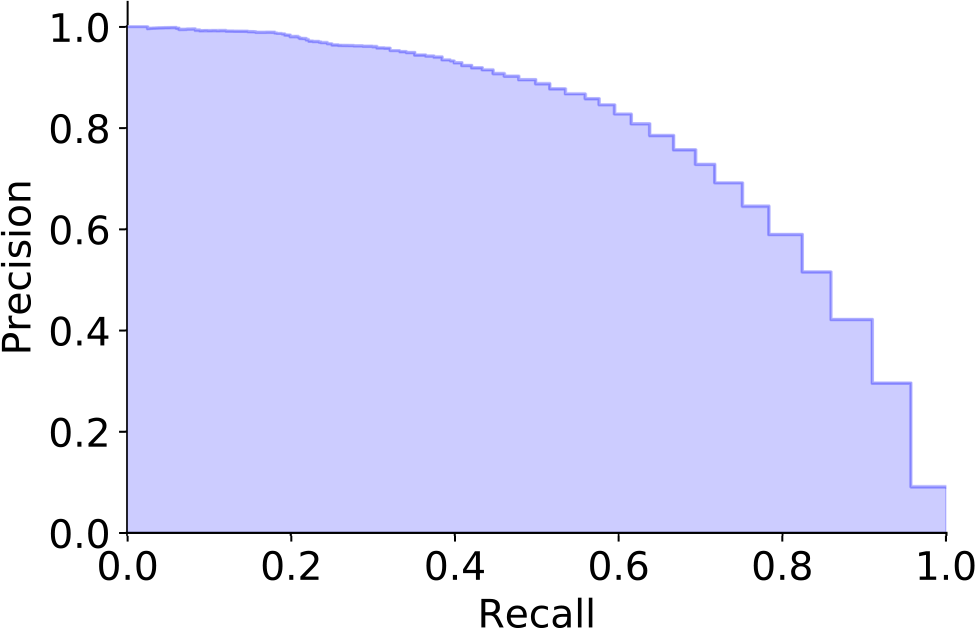
Performance of a training-counts-only classifier: As part of the investigation of the impact of a protein’s frequency in the training set on classifier performance, we created a naive classifier which simply predicts that a pair interacts with probability corresponding to the minimum number of times either of a pair exists in a positive interaction in the training set, normalized over the maximum number of times any such protein appears. We find that this naive classifier actually performs rather well in within-species cross-validation, achieving an AUPR of 0.784. However, such a classifier would output a probability of 0 for all interactions in a data set which did not share any proteins with the training set, as is the case in the cross-species setting.

#### D-SCRIPT performs better on infrequently occurring proteins

To further investigate the finding above, we ranked proteins in the human PPI network by their frequency of occurrence. For a set of quantiles *q ∈*[0, 1], we evaluated out-of-sample D-SCRIPT and PIPR predictions on the human PPI sub-network consisting only of proteins of rank *q* or lower; here, lower *q* corresponds to a lower frequency of occurrence. In absolute terms, as the table below indicates, both D-SCRIPT and PIPR become more accurate at higher *q*. However, D-SCRIPT has a relative advantage at lower *q* (i.e., infrequently occuring proteins) while PIPR performs better at higher *q*. In other words, PIPR’s better within-species performance can be traced to it being more accurate on proteins that occur frequently. This also suggests an explanation for PIPR’s lower cross-species generalizability than D-SCRIPT: when making predictions on an entirely new set of proteins in a different species, knowing the relative frequencies of proteins in the training data might not be particularly useful.

The difference between D-SCRIPT and PIPR might stem from their respective architectures. The protein representation learned by D-SCRIPT is constrained to be a linear projection of the Bepler & Berger pretrained embedding, albeit with ReLU and dropout layers. This regularizes how much frequency information can be incorporated into the model; we note that the Bepler and Berger model was trained with data on individual proteins and would not reflect PPI frequency information. In contrast, PIPR’s design allows for a lot more leeway in training each protein’s representation. This flexibility may allow PIPR to better incorporate the occurrence frequencies into its representation, helping its within-species performance but potentially hurting its cross-species generalizability.

**Table A.2:** Area under precision-recall curve (AUPR) of PPI prediction performance for interactions where proteins appear rarely in the training set. We created increasingly restrictive subsets of the test data, where no protein in the subset may appear more than some cutoff number of times in the training set. While PIPR outperforms D-SCRIPT when frequently trained on proteins are present, at lower quantile values *q*, D-SCRIPT performs relatively better than PIPR -although the absolute performance of both classifiers drops off as the data set is more challenging.

| Cutoff Quantile ($q$) | Cutoff Value | % of Test Pairs | PIPR | D-SCRIPT |
| --- | --- | --- | --- | --- |
| 1.0 | 209 | 100 | **0.830503** | 0.591353 |
| 0.9 | 12 | 75.31 | **0.206161** | 0.118704 |
| 0.8 | 5 | 61.12 | 0.061128 | **0.068445** |
| 0.7 | 2 | 46.85 | 0.014476 | **0.024299** |
| 0.6 | 1 | 38.03 | 0.005838 | **0.011789** |
| 0.5 | 0 | 25.06 | 0.001373 | **0.006673** |

### A.5 Functional Enrichment Using GOGO Distance as the GO Term Similarity Metric

To compute within-cluster functional similarity in the main text, we took the simple but robust approach of mapping GO terms to a limited set of GO Slim terms and then computing Jaccard similarity across the latter (Section 3). In Figure A.2, we used a recently introduced approach from the literature, Zhao et al.’s GOGO algorithm [42], to compute the similarity between pairs of GO terms; we then aggregate these to get within-cluster similarity scores. Our evaluation here yields similar results as presented in the main text: D-SCRIPT produces clusters with higher functional similarity than PIPR.

### A.6 GO Enrichment of SARS-CoV-2 Proteins

We performed a preliminary study to predict viral-host interactions between SARS-CoV-2 and human proteins wherein we compared the sets of over-represented GO terms for human interactors of SARS-CoV-2 proteins, as predicted by D-SCRIPT or PIPR, with those over-represented in the experimentally-determined human interactors (Gordon et al. [43]). Figure A.3 shows the relative similarity of computationally predicted annotations to the experimentally-determined annotations for each SARS-CoV-2 protein. Overall, we found that sets of enriched terms computed using the D-SCRIPT network overlap slightly more with the true network than those computed using the PIPR network (*p* = 0.059). Among the putative accessory factors (ORF * and Protein 14), D-SCRIPT performs significantly better (mean Jaccard similarity 0.029 vs. 0.118, *p* = 0.022, paired one-tailed *t*-test). Visually, PIPR seems to be somewhat better at predicting interaction partners for the non-structural proteins (NSP *), although D-SCRIPT still has a slightly larger mean similarity (0.183 vs. 0.222, *p* = 0.221). While D-SCRIPT performs better on the intensively studied spike (S) protein, PIPR shows a higher overlap for the nucleocapsid (N). Neither method predicts enriched terms for the other structural proteins encoding the envelope (E) and membrane (M) (0.149 vs. 0.121, *p* = 0.672 across the four proteins).

**Figure A.2:**
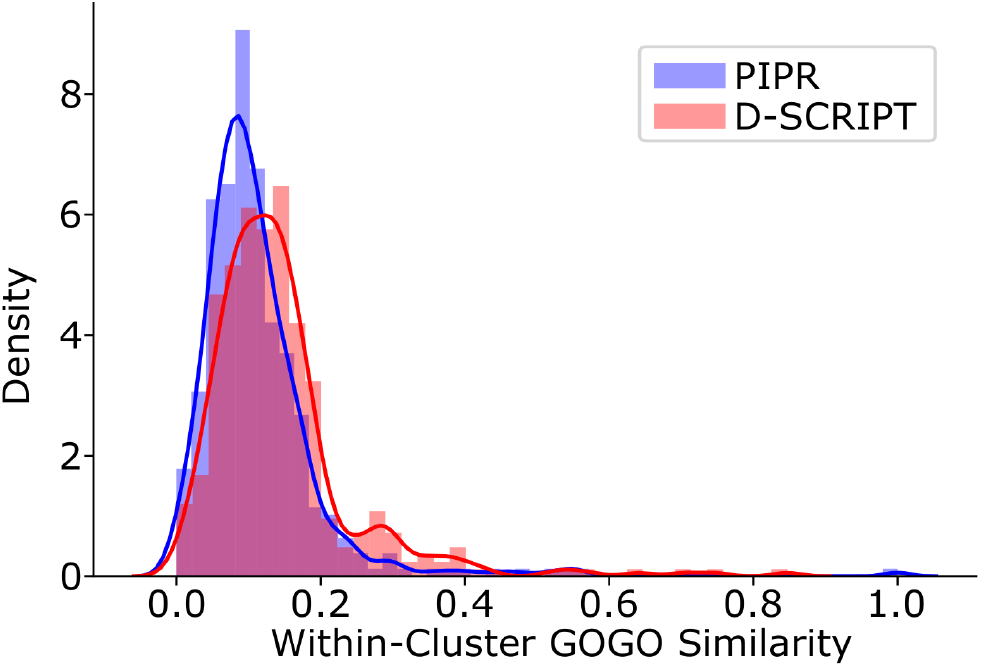
Distribution of within-cluster similarity scores using GOGO Protein Distances. D-SCRIPT recovers significantly more functionally coherent clusters (*p* = 3.94*e −*06) than PIPR when similarity between proteins is measured using GOGO [42]. Clusters of interacting proteins were constructed from the PIPR and D-SCRIPT networks using spectral clustering, where distances between proteins were computed using the diffusion state distance (DSD) metric [38]. We compute the average similarity between pairs of proteins in each cluster. Similarity between proteins is calculated as GOGO Biological Processes similarity using all FlyBase GO term annotations.

### Methods

Candidate pairs were generated using the viral sequences from Gordon et al. [43] and 19,777 human sequences from the STRING database (Section 3), and predicted edges using D-SCRIPT and PIPR. We predicted 3,273 edges using D-SCRIPT and 2,922 edges using PIPR. 332 putative true viral-host interactions were taken from Gordon et al. Human sequences were mapped to UniProt sequences identifiers from [43] with sequence similarity *≥* 95% using BLAST [46], and UniProt identifiers were used to identify a set of Gene Ontology terms for the human interactors of each viral protein. Following [43], we identified over-represented GO terms using the clusterProfiler R package (version 3.14.3) [50] with a 1% false discovery rate (FDR). Over-represented GO terms were mapped to a common set of terms taken from the ChEMBL Drug Target GO Slim Subset [51]. For each viral protein, we computed the Jaccard similarity between the set of GO Slim terms enriched in the putative true network and each of the computationally predicted methods. We computed a paired one-tailed *t*-test to statistically compare the relative similarities of D-SCRIPT and PIPR.

### Availability

Virus-host edges predicted using D-SCRIPT or PIPR are available for use by the community at http://dscript.csail.mit.edu.

**Figure A.3:**
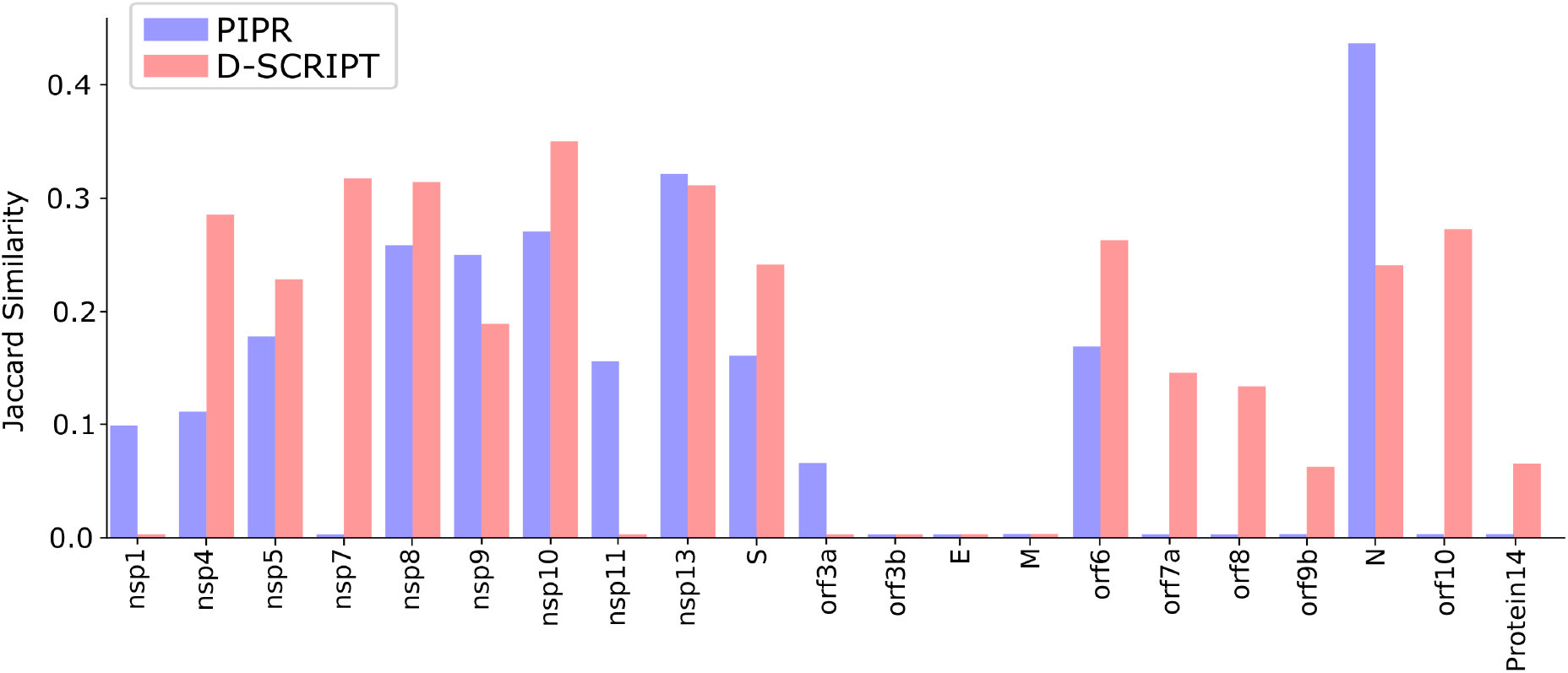
Similarity of Enriched GO Terms to True Network. Interactions between SARS-CoV-2 proteins and human proteins were predicted using D-SCRIPT and PIPR. Viral proteins were then annotated with the enriched ChEMBL GO Slim terms linked to their human interactors. Compared to PIPR, interactions computed with D-SCRIPT show a greater annotation overlap (*p* = 0.059) with those estimated from putative true interactions from Gordon et al. [43]. D-SCRIPT performs especially well at predicting functional enrichments for putative accessory factor proteins, where PIPR recovers none of the enriched functional terms in several cases.

